# A neuroprotective tetrapeptide for treatment of acute traumatic brain injury

**DOI:** 10.1101/2025.04.17.649010

**Authors:** Aman P. Mann, Sazid Hussain, Pablo Scodeller, Hope N.B. Moore, Elan Sherazee, Rachel M. Russo, Erkki Ruoslahti

## Abstract

Traumatic brain injury (TBI) is a major clinical problem because of the high incidence and the severity of the subsequent sequelae. Despite extensive efforts, there are no therapeutic drugs clinically approved for treating acute TBI patients. To address this unmet need, we assessed the activity of the tetrapeptide, CAQK, in mice. When administered intravenously shortly after moderate or severe TBI, CAQK accumulates in the injured brain in mice and pigs. CAQK binds to an extracellular matrix glycoprotein complex that is upregulated in injured brain. Treatment of TBI mice with CAQK resulted in reduction in the size of the injury compared to control mice. There was reduced upregulation of the glycoprotein complex, less apoptosis, and lower expression of inflammatory markers in the injured area, indicating that CAQK alleviates neuroinflammation and the ensuing secondary injury. CAQK treatment also improved functional deficit in TBI mice, with no overt toxicity. Our findings suggest that CAQK may have therapeutic applications in TBI.

## INTRODUCTION

Traumatic brain injury (TBI) is a global public-health problem because of its prevalence and long-term sequelae associated with it. Each year, approximately 70 million individuals worldwide sustain a TBI, which is a leading cause of injury-related death and disability [1]. TBI survivors often suffer from long-term physical disabilities and cognitive disorders, including depression, drug and alcohol abuse, and increased risk of suicide [2]. Current critical care management of TBI focuses on general intensive care support and specific neurocritical care interventions to minimize further damage to the brain by preventing hypoxia, hypotension and hypoventilation. However, there are no clinically proven effective pharmacologic agents to limit secondary injury or enhance repair [3]. Furthermore, advanced imaging and monitoring techniques are not available in smaller trauma care centers, making clinical management of patients with severe TBI even more difficult. Therefore, novel therapeutic approaches are needed to improve clinical outcomes for TBI patients.

TBI is heterogeneous and varies widely in severity, clinical presentation, and pathophysiology [4]. Multiple cascades of signaling events, both acute and delayed, are involved in neuronal cell death following a TBI and play a role in secondary damage. Some of these include blood-brain-barrier disruption, neuroinflammation, oxidative stress, and demyelination. Due to this multifaceted nature of TBI, future therapies to treat TBI patients will likely need a pluripotent mechanism of action to successfully modulate the secondary injury cascade, and, in doing so, maximize the likelihood of a successful clinical outcome. Homing peptides can deliver a variety of therapeutic payloads in high concentrations directly to the site of injury, minimizing systemic side effects, offering an option to facilitate such treatments.

An unbiased screening approach utilizing *in vivo* phage display in a mouse model of unilateral penetrating brain injury revealed a tetrapeptide, CAQK (cysteine-alanine-glutamine-lysine), that showed specific localization into brain injury from intravenous (i.v) administration [5]. Initial data indicated that the peptide extravasates through compromised blood-brain-barrier associated with moderate and severe TBI and does not accumulate in normal brains. In the injured brain, the peptide targets an extracellular matrix (ECM) glycoprotein complex, which is produced in increased amounts and may also be structurally altered in TBI [6]. CAQK binding to injured regions from human brain tissue was also shown [5]. Subsequent studies demonstrated that the injury-targeting capability of CAQK was useful for site-specific delivery of different types of payloads to brain and spinal cord injuries [7–9]. Other disease-specific homing peptides discovered using similar phage library screening have shown inherent biological activity in the absence of a payload that can be therapeutic [10, 11]. We used a pre-clinical controlled cortical impact (CCI) model of TBI to determine whether CAQK might have such an intrinsic activity.

## MATERIALS AND METHODS

### Controlled cortical impact model

All experiments for the CCI model were conducted under an approved protocol (AUF#19-077) of the Institutional Animal Care and Use Committee of Sanford Burnham Prebys Medical Discovery Institute and all experiments were performed in accordance with relevant guidelines and regulations. The CCI model was used as described [12]. Eight- to ten-week-old male C57BL/6 mice were anesthetized with 4% isoflurane (Aerrane; Baxter, UK) in 70% N_2_O and 30% O_2_ and positioned in a stereotaxic frame. Using a head restraint, a 5-mm craniotomy was made using a portable drill and a trephine over the right parietotemporal cortex and the bone flap was removed [13]. Mice were subjected to CCI using the benchmark stereotaxic impactor (Impact One^TM^; myNeuroLab.com) with the actuator part mounted directly on the stereotaxic instrument. The impactor 3 mm tip accelerated down to the 1.0 mm distance, reaching the preset velocity of 3 m/s, and the applied electromagnetic force remained there for the dwell time of 85 ms, and then retracted automatically. The contact sensor indicated the exact point of contact for reproducible results. Face mask anesthesia (1-2% isoflurane in 70%/30% nitrous oxide/oxygen) was used during the entire procedure. In both cases, afterwards, the scalp was closed with sutures, anesthesia discontinued, and mice were administered buprenorphine i.p. for pain control. For the first 2 h post-CCI, mice were closely monitored in their cages.

### Peptide synthesis

The peptides were synthesized on a microwave-assisted automated peptide synthesizer (Liberty; CEM, Matthews, NC) following Fmoc/t-Bu (Fmoc:Fluorenyl methoxy carbonyl, t-Bu: tertiary-butyl) strategy on rink amide resin with HBTU (*N,N,N′,N′*-Tetramethyl-*O-*(1*H*-benzotriazol-1-yl)uranium hexafluorophosphate (OR) *O*-(Benzotriazol-1-yl)-*N,N,N′,N′*-tetramethyluronium hexafluorophosphate)activator, collidine activator base and 5% piperazine for deprotection. Fluorescein tag was incorporated during synthesis at the N-terminus of the sequence for homing studies. Cleavage using a 95% TFA Trifluoro acetic acid followed by purification gave peptides with >90% purity. Peptides were lyophilized and stored at -20°C.

### Affinity Chromatography and Proteomics

For identifying CAQK binding proteins, mouse brains with PBI were collected 6 hrs post-injury. Using liquid nitrogen, the brains were crushed and ground into powder in a mortar and pestle. Next, brain tissue was lysed in PBS containing 200 mM n-octyl-beta-D-glucopyranoside and protease inhibitor cocktail (Roche) as described previously [14] with slight modifications. The clarified lysates were loaded on to CAQK-coated Sulfolink-agarose beads (Pierce) and incubated at 4°C for 3-4 hrs. The column was washed with wash buffer followed by washing with 0.5 mM CGGK control peptide to remove nonspecifically bound proteins. The bound proteins were eluted with 2 mM free CAQK peptide. The eluted factions were pooled, their protein concentration determined by using bicinchoninic acid (BCA) protein assay (Pierce) and the samples were digested using the Filter-aided Sample Preparation (FASP) method [15]. Finally, the digested samples were dried, desalted and subjected to LC-MS/MS analysis at the Sanford Burnham Prebys Medical Discovery Institute’s Proteomics Core facility. All mass spectra were analyzed with MaxQuant software version 1.5.0.25. The MS/MS spectra were searched against the *Mus musculus* Uniprot protein sequence database (version July 2014).

### Proteomics analysis to identify CAQK binding receptor

Serum-free conditioned medium enriched in proteoglycans was obtained by growing U251 cells in phenol-red free DMEM culture media. The collected media was centrifuged to remove cell debris. The supernatant was run on anion exchange diethyl-aminoethyl groups (DEAE) column to capture the proteoglycans. The column was washed with 50 mM Tris-Cl, pH 8.0 and 250 mM NaCl. The purified protein was eluted with two different elution buffers. The elution buffer 1 consisted - 8 M Urea, 50 mM Tris-Cl, 150 mM NaCl, and elution buffer 2 consisted - 50 mM Tris-Cl, 1 M NaCl. The eluted fractions (0.5 mL) were dialyzed in buffer containing 50 mM Tris-Cl, 100 mM NaCl, pH 7.4. The eluted and dialyzed fractions from the DEAE column were subjected to LC-MS/MS analysis at the Proteomics Core facility at Sanford Burnham Prebys Medical Discovery Institute. Chicken chondroitin sulfate proteoglycans (CSPG) mixture (Millipore sigma cat #CC117) was digested with Chondroitinase ABC enzyme (AMS Bio, Cambridge, MA) and subjected to LC-MS/MS analysis to identify the binding partner of CAQK peptide.

### Fluorescence polarization assay

Fluorescence polarization (FP)–based assay was used for solution-based binding of CAQK to different proteins. FP assay was initially set up in 50 μL well final volumes in 96 well plates (Corning Life Sciences), and measurements were carried out using PheraStar FS plate reader (BMG LABTECH, Ortenberg, Germany) in triplicate. FAM-Peptide and protein stock solutions were diluted to desired concentration with assay buffer (10 mM HEPES pH 7.4, 1mM CaCl2, 150 mM NaCl, 0.1% Pluronic). FAM-labeled peptides were protected from light during the experiment to avoid photobleaching. In the binding assay, the concentration of FAM-peptide was kept constant, and the FP was measured over the range of target protein concentrations. The specificity of the binding assay was confirmed by the displacement assay, in which FAM-labeled peptide was competed off from the target protein by unlabeled peptide.

### Cell culture and treatment

U251 cells (from ATCC, Manassas, VA, USA) were cultured at 37°C in 5% CO2 in high glucose DMEM and 10% fetal bovine serum supplemented with penicillin (100 U/ml) and streptomycin (100µg/ml). Conditioned media were obtained by conditioning the growth media for 24 h with confluent cells immediately prior to passaging. CM were centrifuged at 1,500 × g for 10 min and mixed with fresh growth medium at a 1:1 ratio prior to use.

### Homing studies and Immunofluorescence

Animals were intravenously (i.v.) injected, six hours after brain injury, with 50nmoles of FAM-labeled peptide dissolved in PBS and allowed to circulate for 30 minutes. Mice were perfused intracardially with saline, and brains were isolated and imaged using the Illumatool Bright Light System LT-9900 (Lightools Research). For fixing the biological material, brains and organs were fixed in paraformaldehyde (4%) at pH 7.4 for overnight and dehydrated in sucrose gradient (15%-30%) followed by cryosectioning. Sections were permeabilized using PBS-Triton, blocking was carried out using 5% blocking buffer: 5% BSA, 1% goat serum (Jackson Immuno Research), 1% donkey serum (Jackson Immuno Research) in PBS-T. Primary antibodies were incubated in diluted (1%) blocking buffer overnight at dilutions 1/100 or 1/200 at 4C, washed with PBS-T and incubated with secondary antibodies diluted 1/200 or 1/500 in 1% diluted buffer for one hour at room temperature, subsequently washed with PBS-T, counterstained with DAPI 1ug/mL in PBS for five minutes, washed with PBS, and mounted using mounting media, and imaged the same day or the day after using Zeiss LSM-710 confocal microscope. Antibodies used: Human anti-tenascin C (R&D Systems), rat anti-tenascin-C (R&D Systems), Rabbit anti-fluorescein/Oregon Green (ThermoFisher), GFAP antibody 2.2B10 (ThermoFisher). Sections were imaged using a Zeiss LSM 710 confocal microscope. Quantification of immunoreactivity was done in a blinded manner by calculating fluorescent intensity in six randomly selected fields of view and processed using Image J software (v1.8.0, NIH, USA).

### Pharmacokinetic analysis

CAQK (12.5mg/kg) was injected i.v. into mice with CCI at 4 hours post injury. Blood was collected from these mice at different time points and plasma was processed and analyzed by LC/MS after a reduction and alkylation step under reducing conditions to capture total peptide (free peptide and peptide bound to plasma proteins by disulfide formation). Plasma concentration was plotted using WinNonlin. Similarly, for analysis in rats, CAQK (10 mg/kg) was injected i.v. into healthy Sprague-Dawley rats once daily for 7 days. Blood was collected at different time points at Day 1 and Day 7 and plasma was analyzed by LC/MS to analyze CAQK. Plasma concentration of CAQK was plotted. N=3 per time point.

### Peptide treatment studies

For treatment studies, mice with CCI were randomized into two groups (n=6/group) and injected with CAQK (200nmoles dissolved in 100µl PBS) or vehicle (PBS) alone starting at 6 hours after injury. The mice received the same treatment daily until day 7 after injury. At day 10, the mice brains were extracted and placed in 4% paraformaldehyde (PFA) at pH 7.4 overnight, washed with PBS and placed in graded sucrose solutions overnight before optimal cutting temperature compound (OCT) embedding. Ten-micron thick sections covering the entire injury were cut and analyzed by immunofluorescence. To measure the lesion volume, the sections were stained with Nissl stain and the lesion area for each distance from bregma was measured using Imagescope software (Leica Biosystems). Lesion area was plotted against bregma distance and integrated over the entire injury length to obtain the lesion volume using polynomial fit in Origin 2022b.

### Behavioral Testing

Mice with TBI were treated and compared with naïve mice using the following three behavioral tests. Novel object recognition test assayed recognition memory while leaving the spatial location of the objects intact [16]. Mice were individually habituated to a 51cm x 51cm x 39cm open field for 5 min. Mice were tested with two identical objects placed in the field (two 250 ml amber bottles). Each individual animal was allowed to explore for 5 min, now with the objects present. After three such trials (each separated by 1 minute in a holding cage), the mouse was tested in which a novel object replaces one of the familiar objects. Behavior was video recorded and then scored for contacts (touching with nose or nose pointing at object and within 0.5 cm of object). Recognition indexes were calculated using the following formula: # contacts during test/ (# contacts in last familiarization trial + # contacts during test). Rotarod balancing requires a variety of proprioceptive, vestibular, and fine-tuned motor abilities as well as motor learning capabilities [17]. A Roto-rod Series 8 apparatus (IITC Life Sciences, Woodland Hills, CA) was used which records test results when the animal drops onto the individual sensing platforms below the rotating rod. An accelerating test strategy was used whereby the rod started at 0 rpm and then accelerated at 10 rpm. The mice were trained in 3 sets of 3 consecutive trials. Five days later the mice were retested in 3 consecutive trials. The hanging wire test allowed for the assessment of grip strength and motor coordination [18] [19]. Mice were held so that only their forelimbs contacted an elevated metal bar (2 mm diameter, 45 cm long, 37 cm above the floor) held parallel to the table by a large ring stand and were let go to hang. Each mouse was score in three trials separated by 30 seconds. Latency to falling off was also measured up to a maximum of 30 seconds.

### Swine injury studies

This study was approved by the Institutional Animal Care and Use Committee at David Grant USAF Medical Center, Travis Air Force Base, CA (Protocol Number - FDG20210026A). All animal care and use were in compliance with the Guide for the Care and Use of Laboratory Animals in a facility accredited by AAALAC. Castrated male Yorkshire cross swine were acclimated in the facility for at least 10 days before use. They were fed a standard diet and observed for any health problems. Food was withheld the night before surgery, but the pigs had access to drinking water. They were premedicated, intubated, and anesthetized by a trained veterinarian. Premedication and anesthesia induction was achieved with Tiletamine/Zolazepam (Telazol) 6.6-8.8 mg/kg by intramuscular injection the endotracheally intubated and mechanically ventilated. Animals were positioned supine followed by placement of a cardiac monitor, temperature probe, end tidal carbon dioxide monitor, and oxygen saturation monitor. Ventilator settings were adjusted to maintain end tidal carbon dioxide at 40 ± 5 mm Hg. Ophthalmic ointment was applied to each eye to prevent corneal drying. All animals were kept on a warming blanket set to 39 °C to prevent hypothermia. Additional warm air blowers were used if needed to maintain body temperature at or above 37 °C. Crystalloid fluids were administered at 5 mL/kg/hr throughout the experiment. Venous blood was drawn for a preoperative complete blood count (CBC) and complete metabolic profile (CMP). Animals were excluded from further use if their white blood cell count exceeded 25 x 109/L, they were anemic (PCV < 25%), or if their blood chemistry (blood urea nitrogen, creatinine, alanine transaminase, aspartate transaminase, or total protein) values are elevated beyond reference intervals. The craniotomy and vascular access sites were clipped and cleaned of any gross debris. A femoral artery was cannulated percutaneously for the placement of an arterial line for blood pressure measurements and blood gases. A vascular line was placed in the left external jugular vein for venous blood collection and fluid/drug administration. In the prone position, the skin overlying the frontal region of the skull was incised and the soft tissues dissected to expose the skull. A 21 mm burr hole was created on the right frontal bone, 3 mm lateral to the sagittal suture and 7 mm rostral to the coronal suture, taking care not to lacerate the dura. A 2 mm burr hole was made 10 mm lateral and 10 mm caudal to the bregma for insertion of a brain intracranial pressure monitor. The stereotaxic frame was positioned over the head and secured with stabilizer rods inserted into the skull. The exposed cortex was injured with a controlled cortical impact device (Custom Design and Fabrication, Inc., Richmond, VA) mounted to the stereotaxic frame. The parameters include a 20 mm rounded polymer striking tip, 4 m/sec velocity, 12 mm depth, and 200 μ sec dwell time. The craniotomy was packed with bone wax and the skin was sutured in place with continuous sutures. Serial blood samples were drawn from an artery to perform baseline arterial blood gas analyses, CBC, CMP, viscoelastic testing and to obtain sera for later analyses at 5, 10, 15, 30 minutes and then hourly for 6 hours. Standardized supplemental fluids, norepinephrine infusions, and correction of any electrolyte and glucose abnormalities was performed. A minimum MAP of 65 mm Hg was maintained using 500 mL crystalloid boluses and norepinephrine infusions (8 mg/250 mL normal saline) starting at 0.01 mcg/kg/min, up to 0.2 mcg/kg/min. Goal electrolyte and lactate concentrations were monitored and corrected as needed. One hour after injury 2 animals were intravenous fluorescently labeled CAQK peptide (2.5mg/kg), delivered as a bolus over 10 minutes by intravenous infusion pump. Animals were monitored for 1 hour following the completion of treatment during which time serial blood samples were drawn as described above. Animals were humanely euthanized by barbiturate overdose 1 hour after the completion of treatment/placebo administration. Perfusion with 3L phosphate buffered saline was performed by cannulating both carotid arteries. The brains were extracted, chilled at -80 °C for 30 minutes, and sliced. Tissue from the heart, lung, liver, intestine, kidney, spinal cord, and adductor muscle was collected and fixed in formalin. The brain was sliced through the injury impact area in 5 mm coronal slices using a brain block (Zivic Instruments, Inc., Pittsburgh, PA). Brain slices were immersion-fixed in 4% (w/v) paraformaldehyde/PBS for 24 hours. Brain slices were transferred to fresh 4% paraformaldehyde for 48 to 72 hours at 4 °C with gentle shaking and then transferred to PBS for 24 hours at 4 °C, followed by dehydration in sucrose gradient (15%-30%) followed by cryosectioning at 10-20 µm for immunohistochemistry.

### Safety analysis

Male C57BL/6 mice (n = 5 per group) with CCI injury were intravenously administrated daily with control PBS or with various concentrations (1mg/kg, 5 mg/kg and 25 mg/kg) of the CAQK peptide. After two weeks, blood was collected from the mice and analyzed for liver and kidney toxicity. Tests performed for liver function (ALP: Alkaline Phosphatase, ALT: Alanine Aminotransferase, TBIL-total bilirubin, GGT- G-glutamyl transferase) and kidney function (BUN: Blood urea nitrogen). Results expressed as mean ± SD. For a more detailed safety analysis, Sprague Dawley rats were randomized to 4 dose groups. CAQK or vehicle (0.9% Sodium Chloride Injection, USP) was administered once daily by i.v. administration to male and female rats at 0 (vehicle),10, 100, or 300 mg/kg/day (Groups 1-4, respectively) for 7 consecutive days. All animals were euthanized and necropsied after blood sample collection for clinical pathology on Day 8. All animals in Groups 1-4 were evaluated for clinical observations, body weights, and clinical pathology (hematology, coagulation, and serum chemistry), gross necropsy observations and major organ weights.

### RNA sample preparation and Data Analysis

Adult male C57BL/6 mice with CCI were treated with CAQK or vehicle (n=3/group) and cortical tissue was microdissected and flash frozen at 14 dpi. RNA was collected, and mRNA copy number was determined using a NanoString nCounter mouse neuroinflammation and neuropathology panels Approximately 75 mg of cortical brain was isolated from sham and TBI mice (for TBI samples, the cortical tissue collected was approximately 5 mm of the cortical area surrounding the ipsilateral injury site). RNA was from isolated using Trizol reagent (Invitrogen, 15,596–018). RNA concentration and quality were assessed using the Nanodrop 2000 spectrophotometer (Thermo Scientific). Total RNA (20 ng/μl) was run on a NanoString nCounter® system for Mouse Neuroinflammation v1.0 panel and Mouse Neuropathology panel (NanoString Technologies, Seattle, WA) to profile RNA transcript counts for over 1400 genes and 20 housekeeping genes. Sample gene transcript counts were normalized prior to downstream analysis and pairwise differential expression analysis was performed with NanoString’s nSolver software Version 4.0. Subsets of genes displayed as heatmaps were normalized across samples as z- scores and then averaged to a single value per group before plotting with GraphPad Prism. Pathway analyses was performed using Rosalind software (https://rosalind.onramp.bio/) developed by OnRamp BioInformatics, Inc. (San Diego, CA).

### Human tissue sections

Human brain tissues with acute traumatic brain injury were obtained from Banner Sun Health Research Institute in Sun City, Arizona. Trauma 1 is a 91-year-old patient with subacute head injury, occipital scalp contusions and subgaleal blood clots, occipital and right temporal subdural hematoma and hemorrhagic contusions of the right anterior temporal pole. Trauma 2 is a patient with moderate TBI who died at age 72 in an automobile accident. The control case was obtained from Brain Tissue Repository maintained by the Center for Neuroscience & Regenerative Medicine (CNRM) at the Uniformed Services University of the Health Sciences (USU) in Bethesda, MD. Control case is from a 63-year-old male without any neurologic diagnosis or any signs of TBI on detailed neuropathologic evaluation. All sections are from the prefrontal cortex region.

### Statistical Analysis

Data were tested for normality and were found to be normally distributed. Accordingly, data are presented as the mean ± SEM unless otherwise noted. Statistical differences between groups were assessed using paired and unpaired Student t test where appropriate All the significance analysis was done using Statistica 8.0 software, using one-way ANOVA or two-tailed heteroscedastic Student’s t test. Values of p<0.05 indicate statistical significance. The details of the statistical tests carried out are indicated in respective figure legends.

## RESULTS

### CAQK binds tenascin C in the ECM of injured brain

We have previously identified a tetrapeptide, CAQK, that accumulates in the ECM of injured brain. Other homing peptides, in addition to accumulating at their target, have been found to had inherent functional effects at the target tissue [20] [10] [11]. We reasoned that that knowing the target molecule CAQK binds to would be likely to give hints of possible nature of such an activity. Previous work has shown that CAQK binds to a chondroitin sulfate proteoglycan (CSPG)-rich protein complex, the expression of which is elevated following an injury [5]. We confirmed this result by testing the binding of CAQK to CSPG complex isolated from chicken brain. Fluorescence polarization assay, which is widely used to measure interactions of small fluorescent ligands with larger binding partners [21], showed dose-dependent binding of fluorescein (FAM)-CAQK to the CSPG complex not seen with a control peptide (Fig. 1A). Pretreatment of CSPG with chondroitinase ABC to remove chondroitin sulfate and dermatan sulfate side chains of proteoglycans had no effect on the FAM-CAQK binding (Fig. EV1A). This suggests that CAQK interacts with a protein, not glycosaminoglycan, in the CSPG complex. To identify a cell line that could be used as a source of the CSPG complex, we tested human glioblastoma cell line U251. FAM-CAQK bound to proteins in the supernatant culture media of the U251 cells and to CSPGs complex purified from these cells (Fig. EV1B). Proteomics analysis showed that tenascin C (TnC) was present in both the U251 and chicken brain CSPG complexes (Fig. EV1C). Indeed, CAQK efficiently and dose-dependently bound to TnC purified from U251 cells in the fluorescent polarization assay (Fig. 1B). Only marginal binding of CAQK to other related glycoproteins, including tenascin R was observed. The cysteine in the peptide was required for the binding, as AAQK peptide or a cyclic, disulfide-bonded peptide containing the CAQK sequence did not show any binding (Fig. EV1D). Furthermore, addition of glutathione or iodoacetamide (to block the free sulfhydryl in CAQK) completely abrogated the binding of CAQK peptide to TnC (Fig. EV1E). Previously, we have shown that i.v. injected CAQK accumulates in injured side of the brain and not in the contralateral side [5]. No other organs showed peptide accumulation except the kidneys, the main site of clearance of small peptides. We also showed that CAQK is specific for brain injuries, as no CAQK accumulation was detected in perforating injuries inflicted on the liver or skin. The reason may be that brain ECM is different from ECM in other tissues in that it mostly consists of a hyaluronan-CSPG-link protein-tenascin complex [22] [23]. Thus, CAQK appears to be a specific probe for TBI.

**Figure 1.**
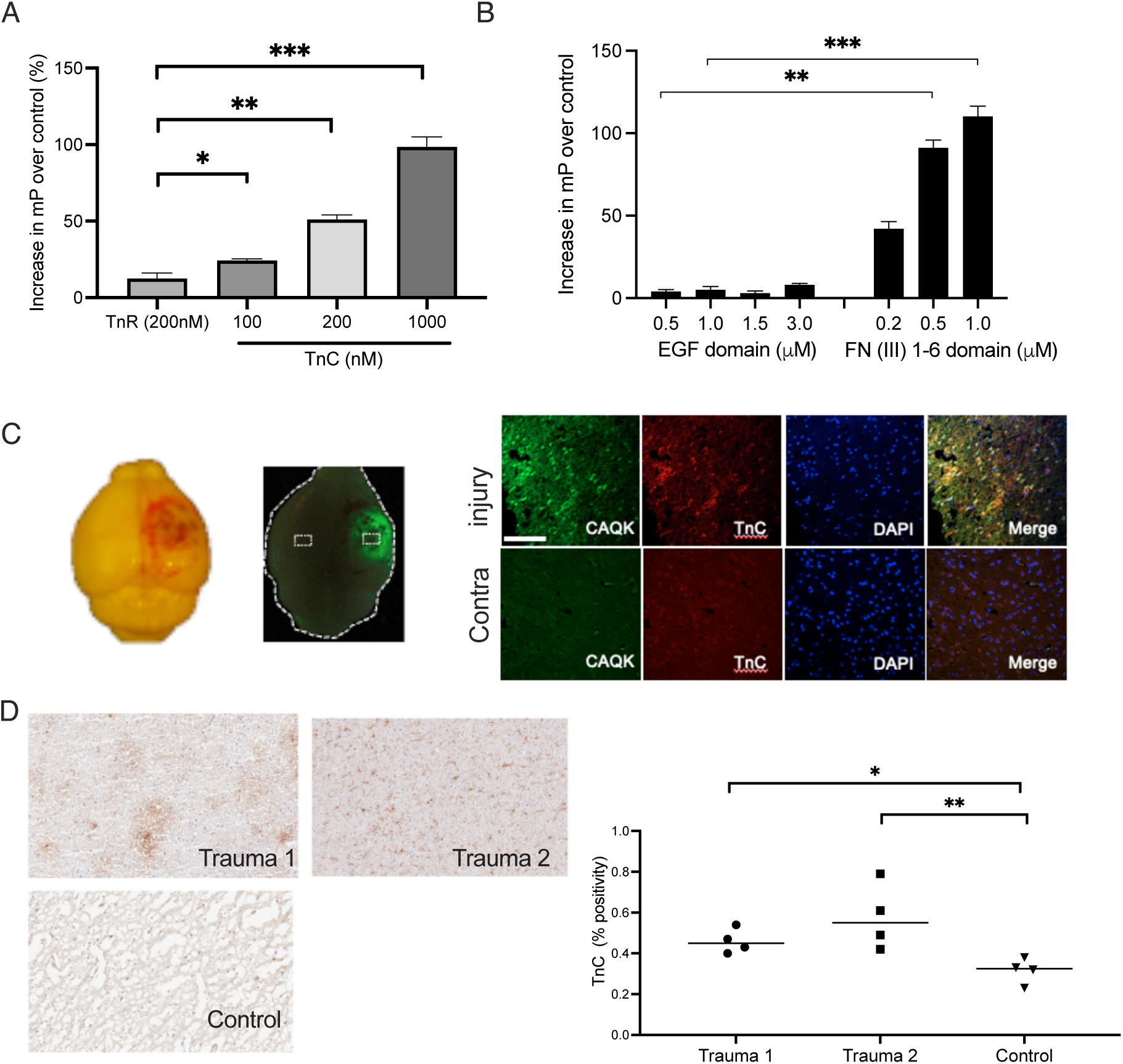
CAQK binds to TnC in the ECM. A. FAM-CAQK binding to purified CSPG complex isolated from chicken brain was analyzed by fluorescence polarization of FAM on the peptide. FAM-CAQK (20nM) was incubated with CSPG complex for 60 minutes at 37°C. Binding was analyzed by measuring the increase in millipolarization units (mP). FAM-AAQK was used as the control peptide. *p < 0.05, **p<0.01 (Fisher’s exact test). All values are mean ± S.E.M from three independent experiments. B. Fluorescence polarization of FAM was analyzed to determine FAM-CAQK (20nM in PBS) binding to purified human TnC after incubation at indicated concentrations for 60 minutes at 37°C. Data expressed as mean ± S.E.M using unpaired t-test. *p<0.05, **p<0.01, ***p<0.005. C. CAQK binding to two recombinant TnC fragments was analyzed by fluorescence polarization of FAM on the peptide. FAM-CAQK (20nM in PBS) was incubated with fragments representing the EGF domains or FN(III) domains of TnC at the indicated concentrations (μM) for 1 hour at 37°C. Data expressed as mean ± S.E.M and analyzed using unpaired t-test. **p<0.01, ***p<0.005. D. To localize the CAQK-binding site more accurately, FAM-CAQK (20nM in PBS) was incubated with a recombinant fragment containing the TnC FN(III) 5-6 domains at indicated concentrations (μM) for 1 hour at 37°C and fluorescence polarization of FAM was analyzed. Cold = excess unlabeled peptide (1μM) added to TnC fragment. Data expressed as mean ± S.E.M. Differences were analyzed using unpaired t-test. *p<0.05, **p<0.01, ***p<0.005. E. Coronal cortical brain sections from mice given CCI and injected with FAM-CAQK i.v. 24 hours after the injury. Left panel shows intrinsic fluorescence of FAM-CAQK seen macroscopically using an Illumatool Bright Light System in the green channel. Higher magnification regions (injured and uninjured) shown in immunostained stained with anti-TnC antibody (red), FAM (CAQK; green) and nuclei (DAPI, blue). Scale bar, 40 μm. F. Immunohistochemical staining for TnC expression on cortical brain sections from frozen brain tissue from human brain tissue from two head trauma patients or a patient with no brain injury as a control.

TnC is an extracellular matrix protein that has been shown to affect the proliferation, migration, survival, and differentiation of cells in the oligodendrocyte lineage [24, 25]. TnC has a multidomain structure consisting of a coiled-coil region followed by multiple EGF-like domains, fibronectin type-III (FN III) domains, and a fibrinogen C-terminal domain [26]. The oligodendrocyte activities have been mapped to the FN (III) repeats in TnC [19]. As homing peptides typically bind to ligand-binding site in proteins [27] [20], we located the CAQK binding site in TnC to see how it relates to the site that affects oligodendrocytes in TBI. Two TnC fragments - one with only the EGF-like domains and other with only the FN III (1-6) domains were expressed from cloned cDNAs. The TnC fragment with the FN III (1-6) domains bound FAM-CAQK in the FP assay, whereas the EGF-like domains did not show binding (Fig. 1C). The minimal fragment we found to be positive for dose dependent FAM-CAQK binding consisted of the FN (III)5 and FN (III)6 domains (Fig. 1D). The TnC binding by CAQK suggested a possible activity in regulating oligodendrocyte responses to injury.

We next tested CAQK targeting and localization in injured brain of mouse model of controlled cortical impact (CCI) injury. CAQK showed specific homing to the injury lesion only (Fig. 1E left panel). TnC has limited expression in the adult brain but is rapidly upregulated in inflammatory processes and injuries to the nervous system [26, 28]. We confirmed high expression of TnC in the injured region of mouse brain using immunofluorescence analysis. TnC expression in TBI coincided with FAM-CAQK homing after intravenous (i.v.) injection (Fig. 1E right panel). In comparison, the contralateral side of the brain showed minimal TnC expression and no peptide homing. We also observed elevated TnC expression in human injured brain tissue compared to normal brain (Fig. 1F) supporting its potential utility for therapeutic application in humans.

### Pharmacokinetic analysis and brain exposure of CAQK

Given the CAQK accumulation in injured brain and binding to TnC in injured ECM, and the functional activities observed with other homing peptides, it seemed possible that CAQK might affect the course of TBI. To prepare for functional testing of CAQK, we performed CAQK half-life analyses using mouse model of CCI. CAQK quantified by mass spectrometry showed biphasic clearance with an initial rapid clearance from the blood (T1/2 = 9 minutes; Fig. 2A). Initial clearance from the circulation through glomerular filtration is expected for a small peptide, explaining the short half-life. A small percentage of the peptide was eliminated with T1/2 = 4.1 hours, likely because of covalent binding of CAQK to plasma proteins, such as albumin, through the free sulfhydryl group ([29] (Fig, EV3). CAQK half-life was also assessed in healthy rats, and it followed a similar biphasic trend as in TBI mice. The half-life remained the same regardless of whether a single injection or daily injections over a week were given (Fig. EV2). At tissue level, CAQK showed significant accumulation (over 8-fold higher) in the injured side of brain than the contralateral side (Fig. 2C). More than 50% of the initially bound peptide remained in the brain after 2 hours with an estimated half-life of about 5.8 hours. We have previously shown that FAM-CAQK is still detectable by fluorescence in the injury area after 3 hours, confirming the prolonged retention at the target [5]. This depot effect, likely due to the binding of CAQK to TnC, suggested that a logistically manageable daily injection schedule could be effective for treatment studies.

**Figure 2.**
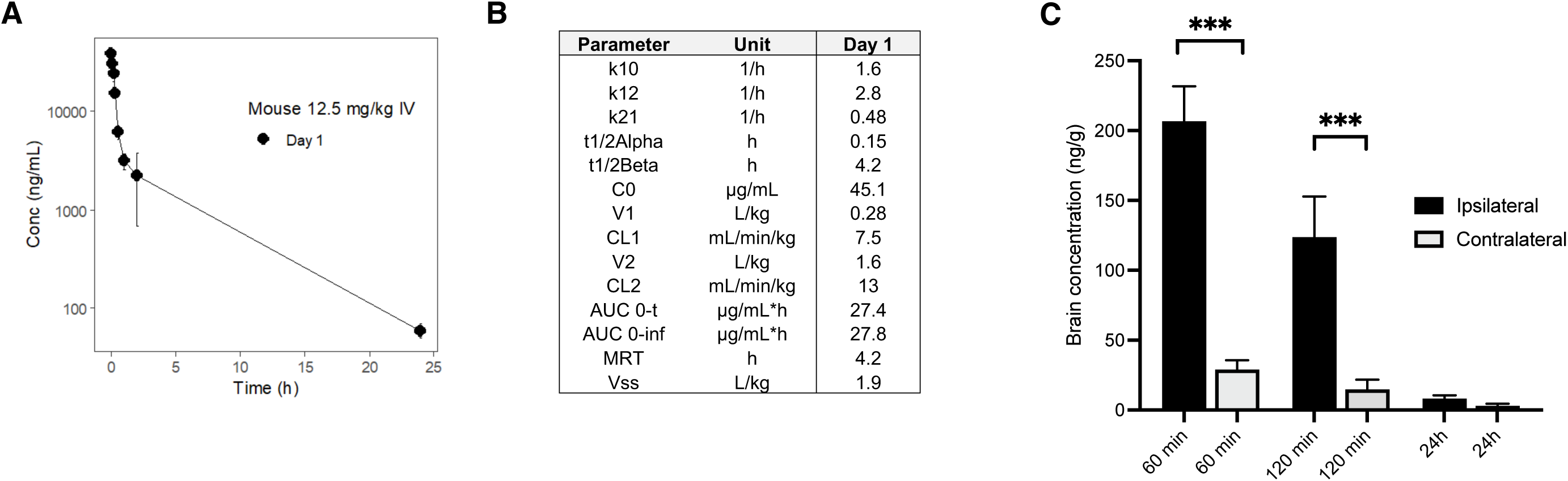
Pharmacokinetic analysis of CAQK. A. Mice with CCI received i.v. bolus of CAQK (12.5mg/kg) at 4 hours after CCI. Blood was collected at different time points and plasma was analyzed by LC/MS. Plasma concentration of CAQK was plotted using WinNonlin. N=3 per time point. B. Plasma clearance data was used to calculate pharmacokinetic parameters using a two compartments analysis. C. CAQK accumulation was analyzed in brain homogenates from injured (ipsilateral) and uninjured (contralateral) side at different time points after i.v. injection and plotted as ng/g of brain tissue. N=3/group. Data were expressed as mean ± S.E.M. Differences were analyzed using unpaired t-test. ***p < 0.03 vs contralateral side.

### CAQK targeting in a gyrencephalic animal model of TBI

Differences in brain anatomy and physiology between rodents and humans has been attributed, in part, to failure to translate TBI therapies promising in mice. To determine whether CAQK targets brain injuries in an experimental animal that has a gyrencephalic brain similar to the human brain (38), we first tested TnC expression in pig brains subjected to CCI. Immunohistochemistry showed robust TnC expression in the brains of CCI pigs not seen in sham injured pigs (Fig. EV3) and i.v.-injected FAM-CAQK homed to the injured area, colocalizing with TnC (Fig. 3). There was minimal or no homing of CAQK to uninjured parts of the brain (Fig. 3) or to sham-injured pig brain, or of a control peptide, or FAM-AAQK, to injured brain (Fig. EV3B). As in mice, the kidneys were the only normal organ positive for FAM-CAQK (Fig. EV3C). The half-life of FAM-CAQK peptide in pig blood was about 30 minutes (Fig. 3C). These results indicate that pigs are an appropriate species for CAQK studies.

**Figure 3.**
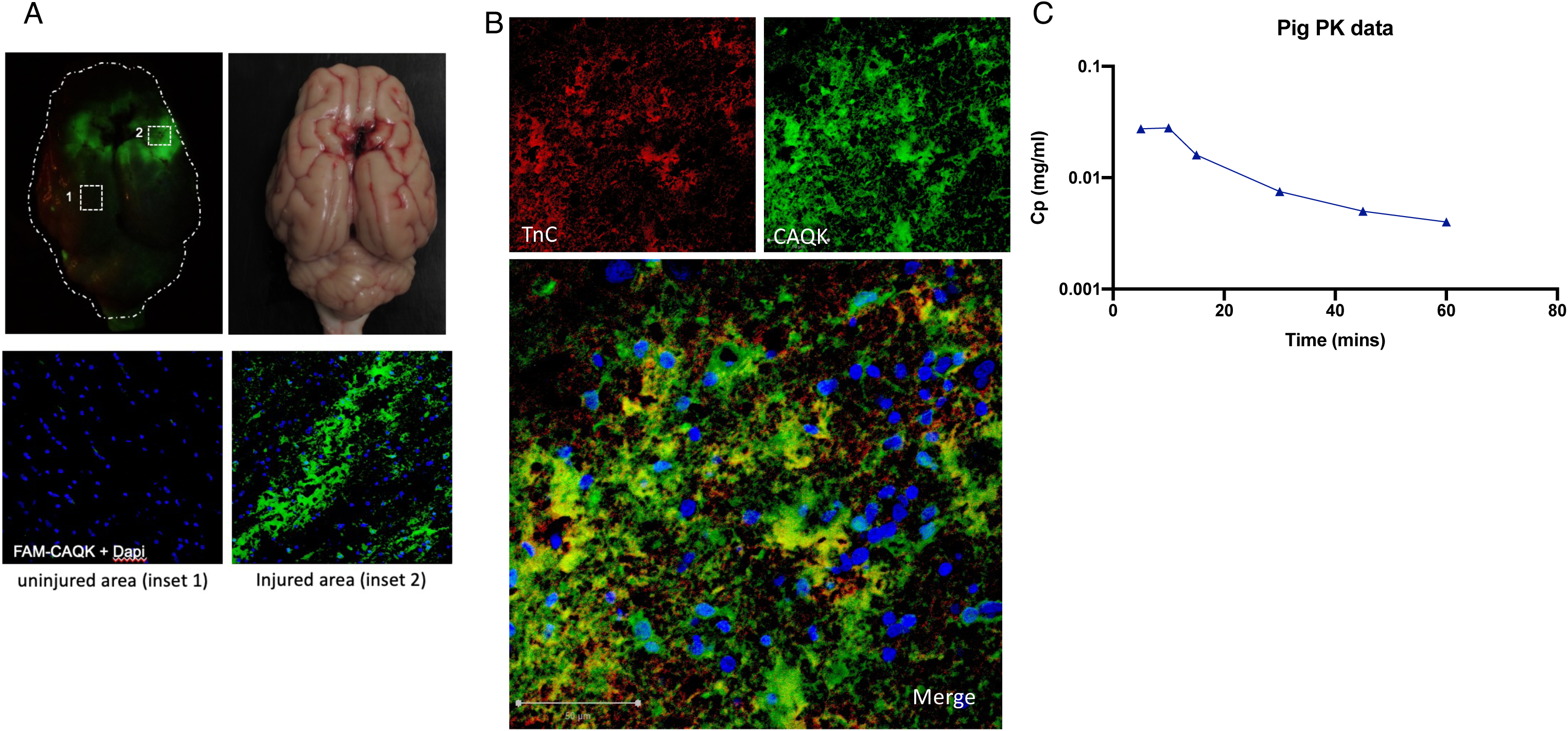
CAQK targets brain injury in pig CCI. A. FAM-CAQK (2.5 mg/kg) was injected i.v. into pigs with CCI at 1-hour post-injury. The peptide was allowed to circulate for 60 minutes. The pigs were perfused, and brains were isolated and imaged using an Illumatool Bright Light System in the green channel. Intrinsic fluorescence of FAM-CAQK is seen macroscopically. Bottom panel shows immunofluorescence staining of FAM signal (green) in uninjured area (*lower left panel*) and injured area (*lower right panel)*. N=2. B. Fluorescence imaging on cortical brain sections from CCI pig injected with FAM-CAQK. Sections were immunostained with anti-TnC antibody (red), FAM (CAQK; green) and counter stained for nuclei with DAPI (blue). Scale bar, 50 μm. C. Plasma concentration (Cp) of FAM-CAQK plotted over time. FAM-CAQK (2.5 mg/kg) was injected i.v. into a pig with CCI at 1-hour post-injury. Plasma concentration was detected by measuring fluorescence from the FAM label in the peptide at different time points.

### Neuroprotective effects of CAQK in TBI

To test CAQK for possible effects on brain injury, mice with CCI were treated with repeated i.v. injections of CAQK (2.5mg/kg per injection) or vehicle control over a period of 7 days post-injury, the time the blood brain barrier, (BBB) remains compromised after injury and enables CAQK entry to the lesion [5]. Macroscopic and histological examination of the brains at day 7 showed greatly reduced (about 50%) tissue loss compared to controls (Fig. 4A-C). CAQK-treated mice also showed significantly reduced apoptosis, as measured by TUNEL staining in the injured areas (Fig. 4D). Neuroinflammation is a major element of the secondary response after TBI. It is associated with reactive gliosis, which is characterized by glial hypertrophy, increased expression of the glial-specific intermediate filament protein GFAP (Glial Fibrillary Acidic Protein), and microglial activation [30] [31]. We observed a drastic reduction in GFAP expression, which reflects the number of activated astrocytes in the lesions and peri-lesion area of CAQK-treated mice at day 7 post-injury (Fig. 4E). There was also a significant reduction in the microglia activation marker, Iba1 (Fig. 4E).

**Figure 4.**
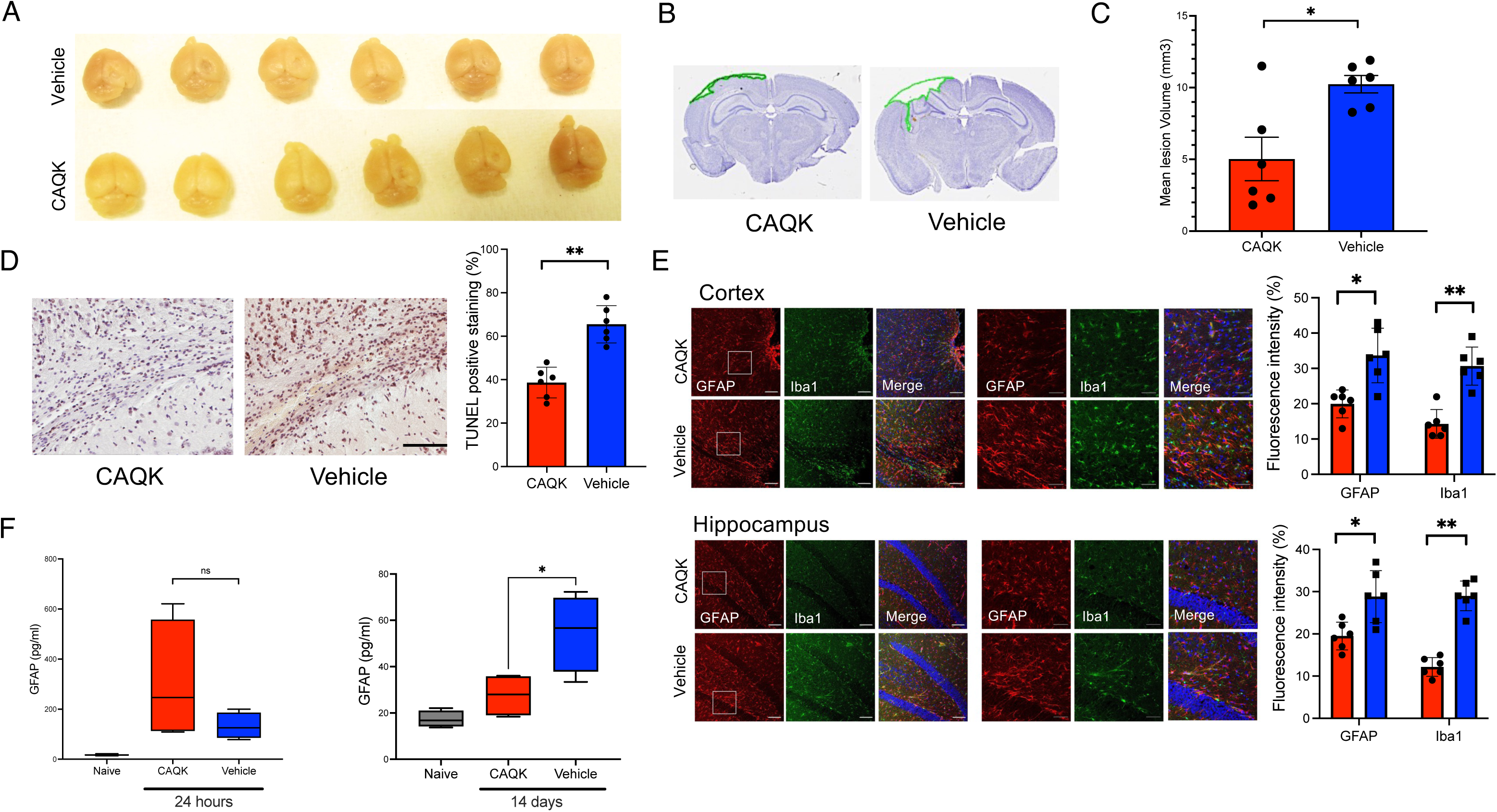
CAQK activity in mouse model of TBI. A. Mouse brains isolated at 7 days post-injury after i.v. treatment with either CAQK peptide (200 nmol (5 mg/kg) per injection) or vehicle (saline) at 6 h after injury and once daily for 7 days. B. Stereological analysis of coronal brain sections (cresyl violet staining) after treatment with CAQK at post-injury day 7 shows significantly reduced tissue loss (n=6/group). C. Quantification of cortical lesion volumes, including the anterior and posterior margins of the contusion, from the stereological analysis in (B). The results indicate a significant difference in lesion areas between the saline-treated and CAQK-treated mice, n = 6/ group. Data were expressed as mean ± S.E.M. Significance was calculated using Fisher’s LSD test (*p < 0.05). Similar results were obtained in three independent experiments. D. Effect of CAQK-treatment on apoptotic cell death examined by TUNEL staining (brown) in an area surrounding commissural fibers of the corpus callosum. Scale bar, 100μm. Quantification of TUNEL staining (% area) analyzed using Image J. Data were expressed as mean ± S.E.M. Differences were analyzed using Fisher’s LSD test (**p < 0.02). E. Effect of treatment on astrogliosis and microglial activation in injured brain. Coronal sections showing cortex and hippocampus from TBI mice treated with CAQK or vehicle were immunostained for GFAP (red) and Iba1 (green). DAPI staining of nuclei is blue. Left panel shows lower magnification. Scale bar, 100μm. Inset were magnified and shown on the right panel. Scale bar, 50μm. Quantification of staining (% area) analyzed using Image J. Data were expressed as mean ± S.E.M. Differences were analyzed using Fisher’s LSD test (**p < 0.02). F. Serum levels of GFAP in TBI mice at 24 hours and 14 days after brain injury. Data compared to naïve mice with no injury and no treatment. Plasma concentration shown (pg/ml; n= 5/group). Data were expressed as mean ± S.E.M. Differences were analyzed using unpaired t-test. *p < 0.05.

GFAP is also a commonly used neurological blood biomarker that has been studied as an indicator of TBI severity [32]. We tested whether GFAP might be suitable a surrogate marker of treatment efficacy. As expected, all mice with TBI showed a dramatic increase in plasma GFAP at 24 hours after injury in all TBI mice. However, at 14 days after injury, the CAQK-treated mice had GFAP levels significantly lower than the vehicle treated mice (Fig. 1F). Another plasma biomarker, NF-L that has been associated with TBI severity, was also tested at 24 hours and 14 days post TBI. Although NF-L levels at day 14 in CAQK treated group trended lower than vehicle group (Appendix Fig. 1), the difference was not statistically significant. Thus, plasma GFAP levels at post-acute phase (day 14) support the accelerated healing of CAQK indicated by the neuroinflammatory markers examined above.

To investigate further the apparent attenuation of neuropathology and neuroinflammation after CAQK treatment, we performed a gene profiling analysis of the TBI lesions. As expected, gene expression associated with neuroinflammation, and neuropathology was robustly increased in TBI mice compared to naïve mice. Volcano plots show the number of neuroinflammation-associated genes that were differentially expressed in vehicle-treated and naïve mice (Fig. 5A, left panel) and CAQK treated and naïve mice (Fig. 5A, right panel). Similar pattern was observed for the neuropathology panel (Appendix Fig. 1). CAQK treatment reduced the number of differentially expressed genes by half relative to the vehicle-treated group (Fig. 5B). A heat map of the highest fold change and statistically significant differential gene expression between CAQK treated vs vehicle treated mice brains is shown in Figure 2C. Pathway analysis showed that the most prominent pathways affected by CAQK were the TYROBP causal network in microglia, and microglial phagocytosis, complement activation and inflammatory response pathways (Fig. 5D). All these pathways are highly correlated with TBI severity and even serve as prognostic indicators [33]. Notably, genes implicated in inflammation and demyelination such as Lipocalin 2, *TGFβ, TNF receptor family, GFAP, MBP* and *S100b* were reduced in CAQK treatment compared to vehicle treated group (Fig. 5C). In addition, genes associated with the complement pathway (such as the C1q gene cluster and C3) which otherwise show prolonged upregulation after TBI, were downregulated in CAQK-treated lesions (Fig. 5E). Interestingly, TnC, the target for CAQK, was also significantly reduced, reaching close to naïve levels following CAQK treatment. These results show that CAQK treatment reverses TBI-related expression of genes associated with inflammation, interferon signaling and neuropathology, reinforcing the other data in building a case for CAQK-mediated improvement of recovery from TBI.

**Figure 5.**
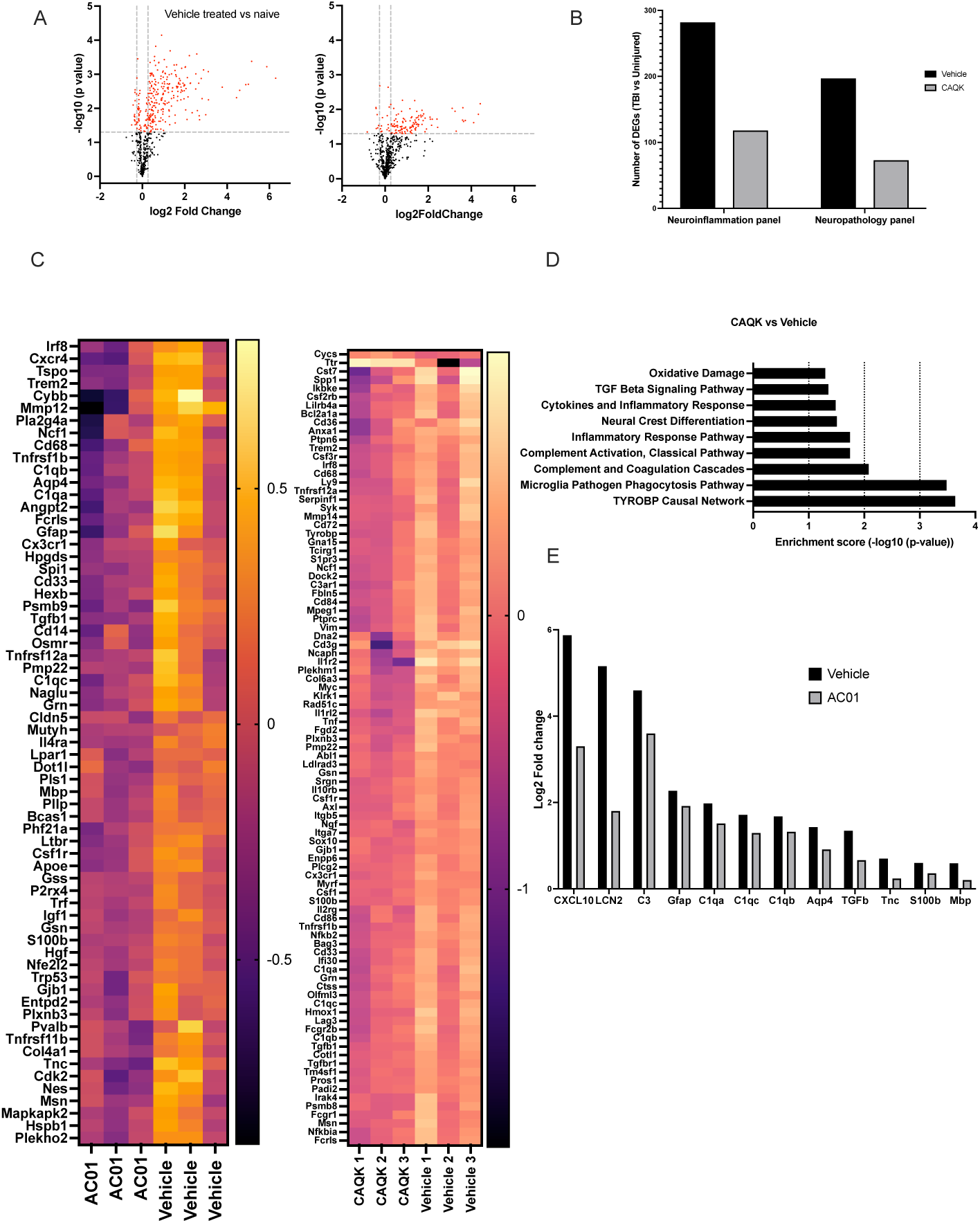
CAQK treatment reduces expression of neuroinflammatory genes in TBI. A. Volcano plots of genes that were expressed differentially at a significant level (p < 0.05) in vehicle treated and naïve mice (left panel) and CAQK treated and naïve mice (right panel) using the neuroinflammation panel (n = 3). B. Number of differentially expressed genes (DEGs) in injured brains at day 14 after injury compared to 12-week-old naïve brains from uninjured mice. Genes were filtered based on fold change > 1.20, p<0.05 as threshold. C. Nanostring based heatmaps showing differential gene patterns in CAQK treated and vehicle treated groups using the neuroinflammation panel (left panel) and neuropathology panels (right panel). Color coding was based on z-score scaling. D. Pathway analysis of the genes that differed most in the neuroinflammation panel between CAQK-treated and vehicle-treated mice after CCI. Differentially expressed genes based on fold change > 1.20, p<0.05 as threshold. E. Analysis of key inflammatory genes implicated in TBI arranged in order of fold change in vehicle treated group. TnC (tenascin C), Gfap (Glial fibrillary acidic protein), Mbp (myelin basic protein), Tgfb1(transforming growth factor-beta 1), S100b (S100 Calcium Binding Protein B).

**Figure 6.**
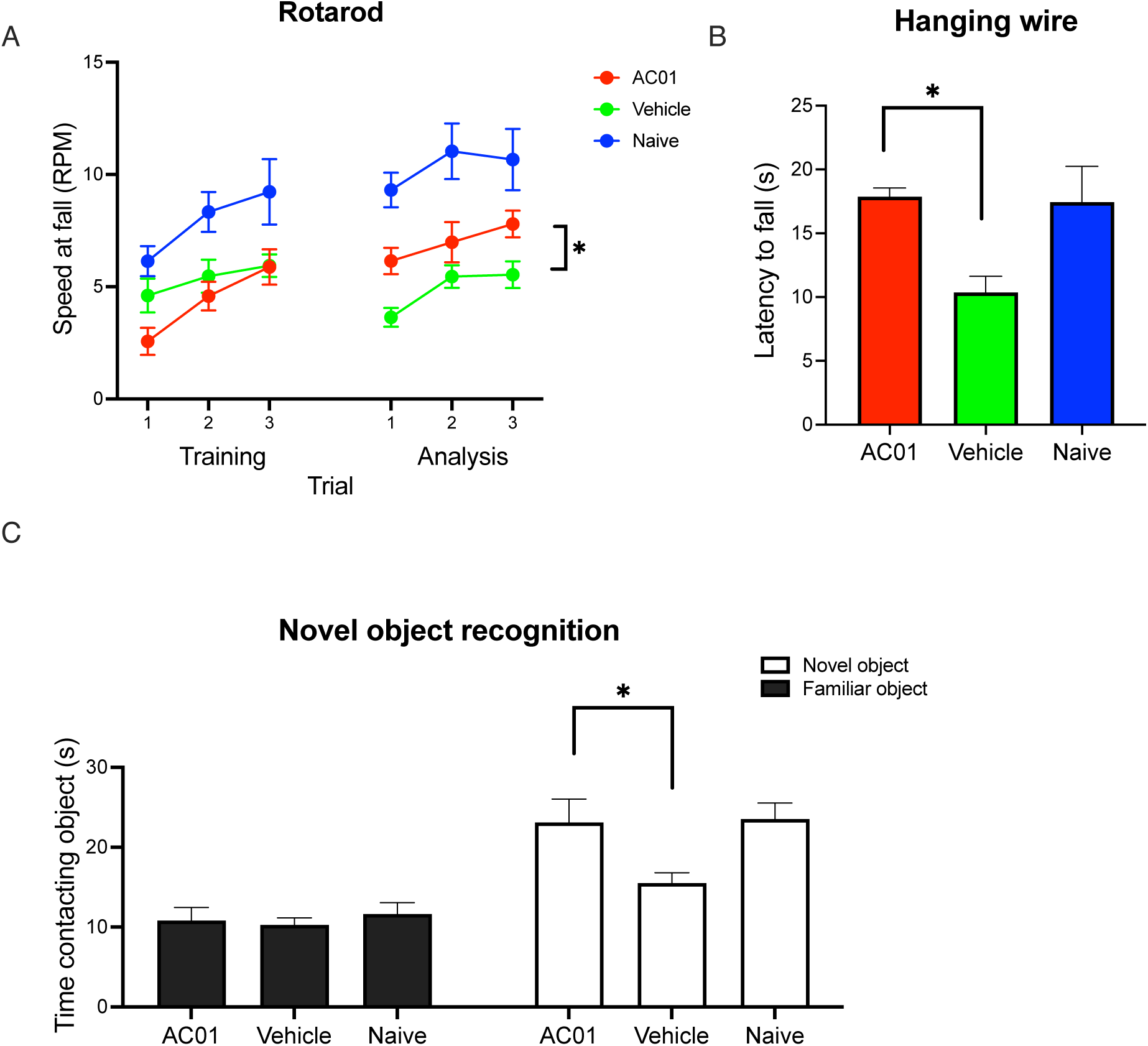
CAQK treatment improves behavioral outcome in TBI mice. A. Rotarod test performed on TBI and naive mice at day 15-18 after injury. Results expressed as mean rotational velocity in rpm at the time of falling off the rod in 3 successive tests. n=10 per group. * p < 0.02, CAQK treated vs. vehicle control (ANOVA). B. Hanging wire test performed in mice on days 22-23 after injury. The latency from the beginning of the test until the time the mouse fell off was timed up to a maximum of 30 s and each animal got 4 trials separated by 30 seconds. n = 10 per group. *p < 0.05. CAQK treated vs. vehicle control. C. Novel object recognition test in mice on days 25-29 after injury. Data is expressed as mean ± SEM. * p < 0.05, one-way ANOVA with Bonferroni test.

The main goal of any treatment of TBI is the best possible functional recovery. An initial “neuroscreen” of TBI mice performed at day 10 showed that the mice had difficulty holding onto the cage top, suggesting the existence of motor problems that would allow comparison of the CAQK-treated and control-treated mice. Reflexes tested by evaluating ear twitch, whisker, and toe pinch responses also suggested signs of recovery of the CAQK-treated mice compared to vehicle treated mice (Appendix Table 1).To assess motor coordination, rotarod and hanging wire tests were conducted on days 15-18 and days 22-23, respectively [18, 19]. In both tests, the CAQK group showed significant recovery compared to vehicle-treated controls (Fig. 3A-B). In a novel object recognition test performed on days 25-29 after TBI, CAQK-treated mice showed greater preference for the novel object than the vehicle treated group (Fig. 3C), also indicating improved recovery of cognitive abilities.

Finally, to assess any potential safety issues, we conducted preliminary safety studies in TBI mice and in healthy rats. The animals were treated daily with different i.v. doses of CAQK up to 300 mg/kg. There were no effects on body weight, organ weight, or clinical pathology (Appendix Tables 2 and 3).

In aggregate, these results show that CAQK treatment results in significant improvement in TBI lesion size and neurological function in mice, suggesting the possibility that CAQK could offer a new treatment option for management of brain injuries.

## DISCUSSION

We describe an inherent neuroprotective role CAQK, a tetrapeptide that was selected to accumulate in areas of brain injury upon intravenous administration and envisioned as a carrier to be used in site-specific delivery of drugs to brain injuries [5]. We show that tenascin C, an ECM component the expression of which is strongly upregulated in brain injury, is the molecule that binds CAQK in TBI lesions. We also describe an inherent neuroprotective role for CAQK. CAQK alone, with no dug attached to it, improves the recovery of mice from acute TBI by histological, biomarker and behavioral criteria. This activity is likely to be a result of the binding of CAQK to TnC. The neuroprotective activity and the demonstration that CAQK also recognizes injuries in gyrencephalic brains in humans and pigs, suggests that this simple and well-defined compound has potential as a new treatment for brain injuries and warrants further study.

Several lines of evidence support the identification of TnC as the target molecule. First, TnC was the only CAQK-binding molecule present in substantial quantities in the two sources we used to prepare ECM. Second, further analysis showed direct binding of FAM-CAQK to recombinant human TnC and to fragments that encompass FN (III) domains 5-6 of TnC. Third, TnC is strongly overexpressed in the ECM of injured regions of the mouse and human brain [5, 34], and CAQK accumulation in brain injuries colocalized with TnC.

TnC, the protein we identify as the CAQK target molecule in brain injury ECM is a regulator of multiple cellular functions [34] suggesting that the TnC binding is likely to be functionally significant beyond making possible the accumulation of CACK at sites of injury. TnC contributes to glial scarring and causes neuroinflammation, blood-brain barrier disruption, and neuronal apoptosis [26]. TnC also inhibits oligodendrocyte differentiation and myelin basic protein (MBP) expression during recovery from a demyelination injury [35],[36] [37]. The underlying signaling cascade requires the binding of the cell surface receptor contactin1 to TnC FN(III) 5 and 6 domains [38]. Our data on CAQK binding to TnC FN(III) 5-6 domains suggests that CAQK could impact these functions of TnC.

Our study shows that CAQK therapy administered within hours after TBI in mice reduced the lesion size, evident both in inspection of the brain and microscopic analysis. The reduction in apoptosis we observed is a likely explanation for the reduced lesion volume. Neuroinflammation was also reduced judging from lower levels of inflammatory markers which included the microglial marker Iba-1 and pathways associated with microglial activation. The strong reduction of Iba-1 is particularly noteworthy because human and animal studies have shown that microglia can be chronically activated for weeks to years after brain trauma [39–42].

TBI initiates a complex cascade of dysregulated pathways that requires a multifunctional therapy. CAQK affected so many different events in the tissue response to TBI that it seems capable of providing the type of pluripotent effect needed to deal with multiple pathways. It may be that CAQK modulates a pathway that is central to the initial response to TBI, and that this hypothetical pathway is governed by TnC.

Importantly, CAQK therapy resulted in improvement in functional recovery of TBI mice. The behavioral effects of CAQK were significant but not as striking as the effects on lesion size and tissue markers. Rodents have relatively small lissencephalic brains in which white matter is less than 20% compared to 80% in large animals and humans, therefore only limited white matter axonal pathology is observed in rodent TBI models [43]. This circumstance has made it difficult to translate mouse results to the clinic. The results reported here show that CAQK also recognizes brain injuries in pigs, a species with a gyrencephalic brain similar as the human brain. We also know from our earlier study [5] that CAQK binds to injured, but not normal, human brain in microscopic overlay assays. Thus, the main prerequisites for efficacy studies in pigs and any subsequent clinical trials are in place.

An important consideration regarding the use of CAQK in treating brain injuries is access to extravascular brain tissue where CAQK binds to the ECM and is retained for a prolonged period of time. The ECM binding may provide a reservoir and favorable localization for CAQK that can potentially amplify its functional effects. Moderate to severe TBI disrupts the structural and physiological integrity of blood vessels resulting in an impaired blood-brain barrier (BBB) that starts as early as 30 minutes after injury, and that can last up to a week after the injury [5, 44], providing a window for systemically administered CAQK to enter the injured target tissue. Disruption of BBB has been observed in TBI patients in the acute phase and given that human brain is not as resilient as mouse brain, the BBB dysfunction may persist well into the chronic postinjury phase [45] [46]. This can provide a much longer treatment window for CAQK than is available in mice. In addition to TBI [47, 48], specific homing of CAQK has been reported in other central nervous system lesions, including, spinal cord injury [7–9], and demyelinating injuries [49]. If the BBB is compromised in these conditions, as it is in TBI, they could potentially also be treated with CAQK.

The tools available for clinicians to treat TBI patients are limited. The current clinical practice in treating TBI patients is essentially limited to supportive care, including the vitally important decompressive craniectomy or other decompression protocols in severe cases of TBI. Cerebrolysin, a peptide preparation generated by controlled digestion of porcine brain proteins is approved for human use in several countries, but has showed only modest improvement of recovery as measured using the Glasgow coma scale [50]. Another deterrent to its use may be the ill-defined nature of the compound. CAQK, in contrast, is a simple, well-defined compound. N-acetyl-L-cysteine (NAC) [51–53], has some similarity with CAQK, as both are small compounds with a free thiol group. The thiol group may contribute to scavenging of tissue-damaging oxygen radicals at the site of injury [54],[55]. The levels of the physiological antioxidant, glutathione (GSH) and its precursors decrease following TBI [56]. The thiol group of GSH is supplied by cysteine. Thus, NAC used to treat liver toxicity caused by acetaminophen [57] could either act as direct ROS scavenger or increase the supply cysteine for the replenishment of GSH [58]. CAQK, with its a free thiol-containing cysteine, may act in a similar fashion.

Given the heterogeneity in presentation and severity of TBI patients, utilization of blood-based protein biomarkers such as GFAP has been approved and validated for initial patient evaluation. GFAP levels correlate with injury severity and are of prognostic value in functional outcome in TBI cases [59]. Our results on TBI mice treated with CAQK show reduced GFAP levels post injury compared to vehicle groups suggesting that GFAP could be used as to monitoring treatment efficacy in future clinical studies. Similarly, multiple studies have shown that serum tenascin-C concentrations of TBI patients are significantly elevated and negatively correlated with favorable outcomes [60, 61]. Our rodent studies show that CAQK treatment results in reduction in TnC expression in injured brain, suggesting that TnC levels might be a potentially useful biomarker.

In conclusion, treatment of TBI mice with CAQK peptide strikingly limits secondary tissue damage from the injury. This, and its apparent lack of toxicity, are encouraging and suggest that further studies toward clinical translation of CAQK are warranted.

## DATA AVAILABILITY

The datasets that were analyzed to support the findings of this study are available from the corresponding author on request.

## ACKNOWLEDGEMENTS

We thank Dr Venkata Ramana Kotamraju at SBP for peptide synthesis, Guillermina Garcia at SBP Core facility for assistance in histology and Alex Campos at the SBP Proteomics facility for proteomics analysis. We also thank Center for Neuroscience & Regenerative Medicine Brain Tissue Repository at the Uniformed Services University of the Health Sciences in Bethesda, MD for sharing human brain tissues. This work was supported by NIH R43NS112050 to A.M and NSF (1548490 and 1660165) to S.H.

