## Supplementary Figures for "A neuroprotective tetrapeptide for treatment of acute traumatic brain injury"

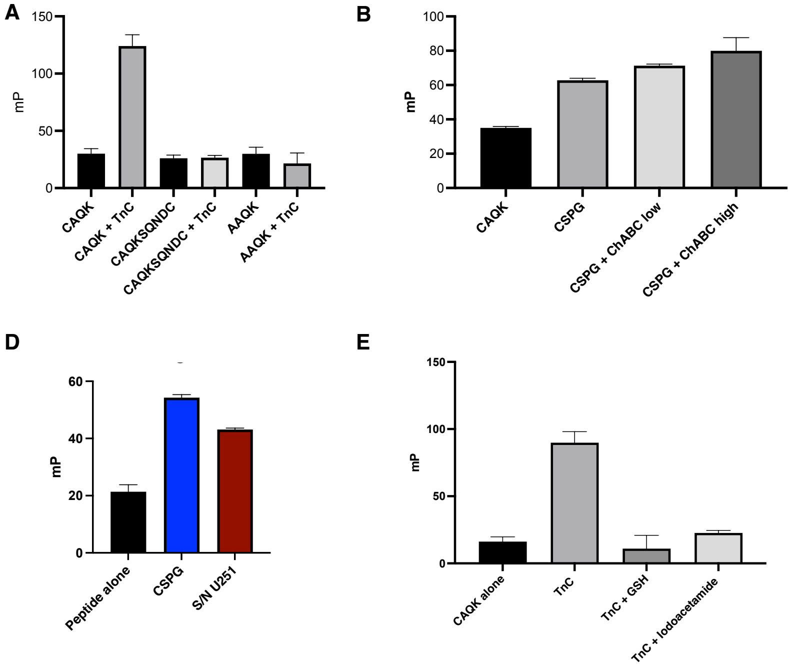

**C**

| Chicken CSPG components | U251 ECM components |
|---|---|
| Neurocan | Laminin subunit alpha-5 |
| Protein-tyrosine-phosphatase | Vimentin |
| Versican core protein | Galactin-3 binding protein |
| Aggrecan | Fibronectin |
| Brevican | Heat shock protein HSP 90-alpha |
| Agrin | Transitional endoplasmic reticulum ATPase |
| Tenascin-R | Tenascin-C |
| Decorin | Nidogen-1 |
| Fibronectin | Filamin-A |
| Tenascin-C | Thrombospondin 1 |

**Figure. EV1. CAQK binding to CSPG components. A.** Fluorescence polarization (FP) measurement of CAQK binding to CSPG. FAM-CAQK (20nM) was incubated with CSPG (200nM) for 60 minutes at 37°C. Binding with CSPG pretreated with chondroitinase ABC (chABC) at two concentrations low (5mU) and high (15mU) was compared to untreated CSPG. **B.** FP assay to assess binding of CAQK to brain ECM from different sources. FAM-labeled CAQK (20nM) was incubated for 1 hour at 37°C with purified CSPG isolated from chicken brain (1□M) or supernatant collected from cultured U251 human glioblastoma cell line. **C.** Top hits from proteomic analysis of CSPG complex isolated from chicken brains and U251 conditioned media **D.** FP assay of binding of different peptides to TnC. FAM-labeled peptides (20nM) were incubated with TnC (1□M) for 1 hour at 37□C. **E**. FP assay to assess the effect of thiol group in cysteine of CAQK to binding to TnC. FAM-labeled CAQK (20nM) was incubated with TnC (1□M) for 1 hour at 37°C in the presence of GSH and Iodoacetamide.

A

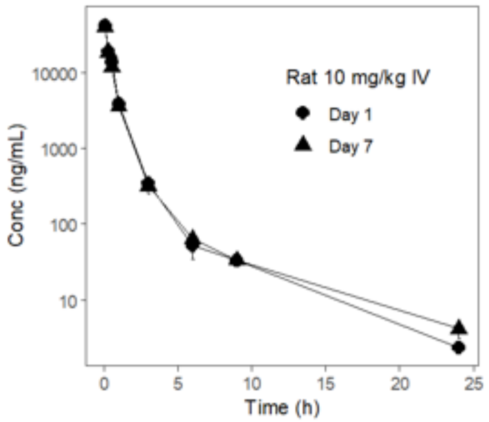

B

| Parameter | Unit | Day 1 | Day 7 |
|---|---|---|---|
| k10 | 1/h | 1.7 | 1.7 |
| k12 | 1/h | 0.074 | 0.090 |
| k21 | 1/h | 0.19 | 0.16 |
| t1/2Alpha | h | 0.39 | 0.38 |
| t1/2Beta | h | 3.9 | 4.6 |
| C0 | µg/mL | 30.8 | 28.9 |
| V1 | L/kg | 0.33 | 0.35 |
| CL1 | mL/min/kg | 9.3 | 10.0 |
| V2 | L/kg | 0.13 | 0.19 |
| CL2 | mL/min/kg | 0.40 | 0.52 |
| AUC 0-t | µg/mL*h | 18.0 | 16.7 |
| AUC 0-inf | µg/mL*h | 18.0 | 16.7 |
| MRT | h | 0.82 | 0.90 |
| Vss | L/kg | 0.45 | 0.54 |

C

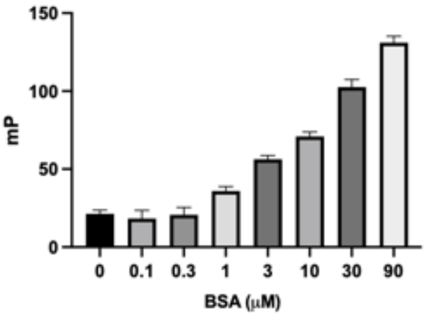

**Figure EV2**. **CAQK plasma clearance in rats. A.** CAQK was injected daily i.v. in healthy Sprague-Dawley rats at 10mg/kg dose for 7 days. Blood was collected at different time points at Day 1 and Day 7 and plasma was analyzed by LC/MS to analyze CAQK. Plasma concentration of CAQK was plotted. N=3 per time point. Plasma clearance data was also used to calculate pharmacokinetic parameters in panel B. **C**. FP measurement of CAQK binding to BSA. FAM-CAQK (20nM) was incubated with BSA at indicated concentration for 1 hour at 37°C.

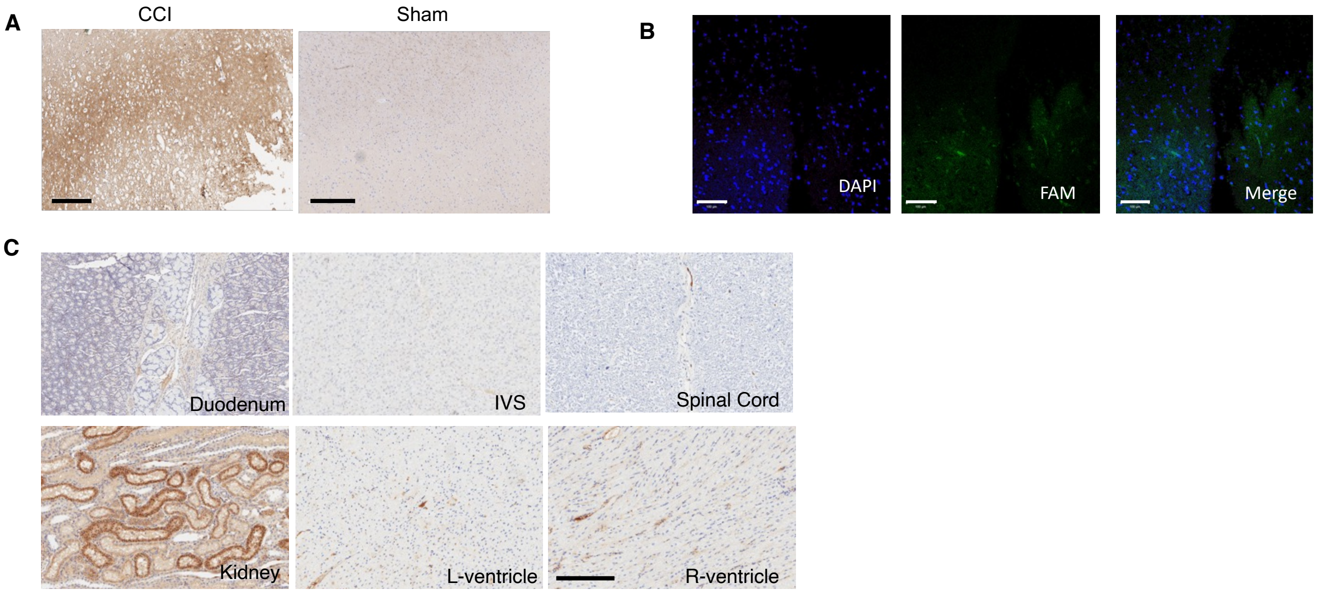

**Figure EV3. Peptide targeting in pig model of TBI. A** TnC expression is upregulated in pig CCI. Immunohistochemical staining for TnC on cortical pig brain sections shows elevated tenascin C expression in the cortex surrounding CCI brain injury compared to the cortex of a sham-injured animal. Scale bar – 300µm. **B.** Control peptide does not accumulate in the brain of pig with CCI. Fluorescence imaging on cortical brain sections from CCI pig injected with FAM-AAQK. Sections were immunostained with anti-FAM (AAQK; green) and nuclei (blue). Scale bar – 100 µm. **C.** CAQK accumulation in different organs in pig CCI**.** Male Yorkshire pig with CCI was injected with FAM-CAQK at 6 hours post-injury. IHC staining of FAM label on fixed paraffin embedded tissue sections from different organs collected 1 hour after peptide injection. Scale bar, 200µm. IVS = interventricular septum

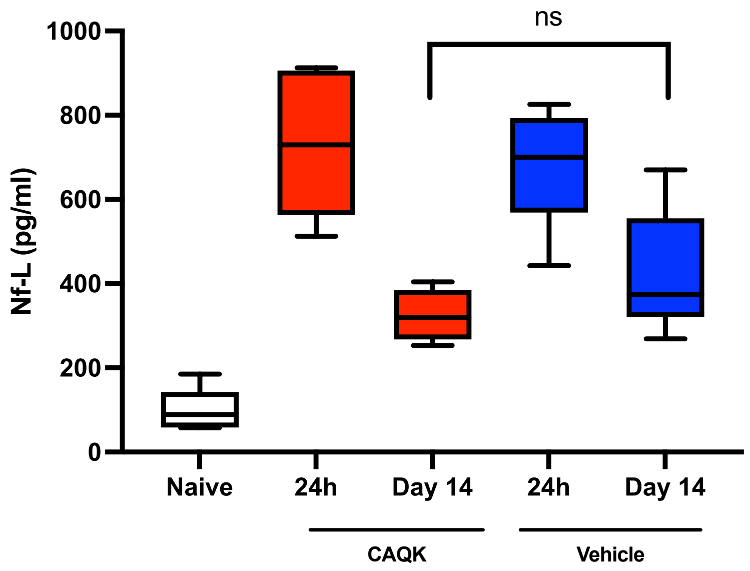

**Appendix Fig. 1. Nf-L levels in TBI mice after treatment.** Serum levels of Nf-L in TBI mice at 24 hours and 14 days after brain injury. Data compared to naïve mice with no injury and no treatment. Plasma concentration shown (pg/ml; n= 5/group). Data were expressed as mean ± S.E.M. Differences were analyzed using unpaired t-test. n.s not significant.

A

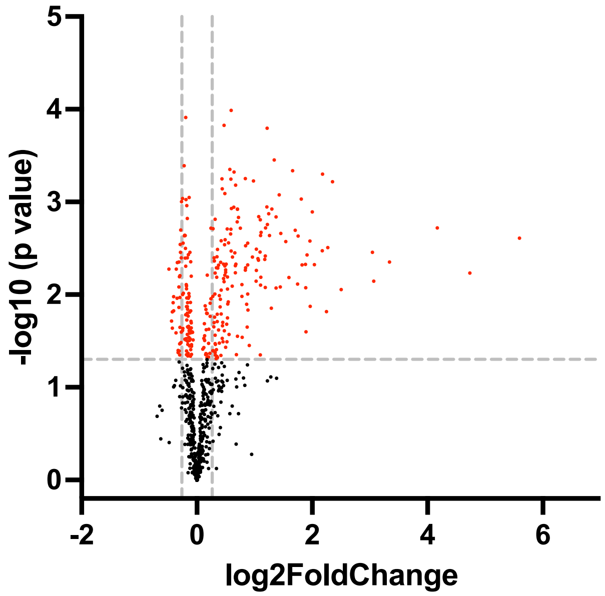

B

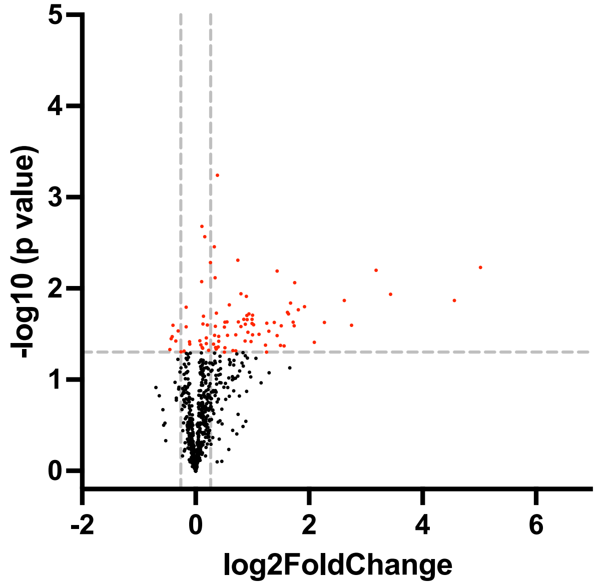

**Appendix Fig. 2.** Volcano plots of significantly differentially expressed genes ($p < 0.05$) compared between vehicle treated and naïve mice (A) and CAQK treated and naïve mice (B) using the neuropathology panel from Nanostring (n = 3).

| | Naïve | Vehicle | CAQK |
|---|---|---|---|
| | N=8 | N=8 | N=8 |
| **Motoric, muscular abilities** | | | |
| Positinal passivity | 0 | 0 | 0 |
| Trunk curl | 0 | 0 | 0 |
| Forepaw reaching | 100.00% | 100.00% | 100.00% |
| Righting reflex | 100.00% | 100.00% | 100.00% |
| Wire hanging (s) | 23.18 | 13.49 | 25.01 |
| **Reflexes** | | | |
| Eye blink | 62.50% | 50.00% | 37.50% |
| Ear twitch | 50.00% | 12.50% | 37.50% |
| Whisker response | 100.00% | 25.00% | 100.00% |
| Toe pinch response | 75.00% | 12.50% | 75.00% |
| **Reactivity** | | | |
| Moving away on petting | 100.00% | 100.00% | 100.00% |
| Struggling on restraint | 100.00% | 100.00% | 100.00% |
| Vocalizing on restraint | 50.00% | 37.50% | 25.00% |
| Dowel biting (3 pt. scale) | 1.13 | 0.75 | 1.13 |
| **Empty cage behaviors** | | | |
| Freezing on transfer | 12.50% | 0 | 12.50% |
| Wild running | 0 | 0 | 0 |
| Stereotypies | 0 | 0 | 0 |
| Cage exploration (3 pt. scale) | 1.63 | 1.25 | 2 |

**Appendix Table 1**. Neuroscreen performed on mice at day 10 after CCI to assess for initial functional deficits. All data are expressed as a percentage of mice that displayed the phenotype shown, unless an alternative measure is stated in parentheses.

| Groups | ALP (U/L) | ALT (U/L) | GGT (U/L) | TBIL (mg/dL) | BUN (mg/bL) |
|---|---|---|---|---|---|
| **1 mg/kg** | 45 ± 5.3 | 24.3 ± 6.8 | < 5 | 0.3 | 21.3 ± 3.5 |
| **5 mg/kg** | 47.4 ± 8.2 | 23.4 ± 4.9 | < 5 | 0.3 | 22.2 ± 1 |
| **25 mg/kg** | 42.6 ± 7.6 | 34.8 ± 13.2 | < 5 | 0.3 | 24.8 ± 2.05 |
| **Vehicle; PBS** | 45.7 ± 3.2 | 35.7 ± 10.8 | < 5 | 0.3 | 21.3 ± 3.1 |

**Appendix Table 2. Safety study in CCI mice.** C57BL/6 mice (n = 5 per group) with CCI injury were intravenously injected daily with vehicle control (PBS) or with the shown concentration of the CAQK peptide. After two weeks, blood was collected from the mice and analyzed for liver and kidney toxicity. Tests performed for liver function: ALP: Alkaline Phosphatase, ALT: Alanine Aminotransferase, TBIL- total bilirubin, GGT- G-glutamyl transferase, and for kidney function: BUN: Blood urea nitrogen. Results expressed as mean ± SD.

**Appendix Table 3. 7-Day IV Dose Study in Rats.** Sprague Dawley rats were randomized to 4 dose groups as shown in the table below. CAQK or vehicle (0.9% Sodium Chloride Injection, USP) was administered once daily by i.v. administration to male at 0 (vehicle),10, 100, or 300 mg/kg/day (Groups 1-4, respectively) for 7 consecutive days. The rats were euthanized and necropsied after blood sample collection for clinical pathology on Day 8.

A. Mean Red Blood Cell and Coagulation Parameters

| Sex: Male | | RBC (M/uL) [g] | HGB (g/dL) [g] | HCT (%) [g] | MCV (fL) [g] | MCH (pg) [g] | MCHC (g/dL) [g] | ABSRET (10^9/L) [g] |
|---|---|---|---|---|---|---|---|---|
| Group 1 - 0 mg/kg | Mean | 7.003 | 14.87 | 43.20 | 61.67 | 21.20 | 34.47 | 297.80 |
| | SD | 0.295 | 0.21 | 3.47 | 2.42 | 0.70 | 2.57 | 95.55 |
| | N | 3 | 3 | 3 | 3 | 3 | 3 | 3 |
| Group 2 - 10 mg/kg | Mean | 7.077 | 14.50 | 44.13 | 62.33 | 20.50 | 32.90 | 374.87 |
| | SD | 0.125 | 0.53 | 1.11 | 0.81 | 0.36 | 0.61 | 43.35 |
| | N | 3 | 3 | 3 | 3 | 3 | 3 | 3 |
| Group 3 - 100 mg/kg | Mean | 7.160 | 14.33 | 43.43 | 60.70 | 20.03 | 33.03 | 354.37 |
| | SD | 0.572 | 0.95 | 2.75 | 1.31 | 0.49 | 0.12 | 9.68 |
| | N | 3 | 3 | 3 | 3 | 3 | 3 | 3 |
| Group 4 - 300 mg/kg | Mean | 7.070 | 14.70 | 42.63 | 60.37 | 21.00 | 34.67 | 345.93 |
| | SD | 0.791 | 0.35 | 3.82 | 2.05 | 2.33 | 2.80 | 22.21 |
| | N | 3 | 3 | 3 | 3 | 3 | 3 | 3 |

B. Mean White Blood Cell Parameters

| Sex: Male | | WBC (K/uL) [g] | ABNEUT (K/uL) [g] | ABLYMP (K/uL) [g] | ABMONO (K/uL) [g] | ABEOS (K/uL) [g] | ABBAS (K/uL) [g1] |
|---|---|---|---|---|---|---|---|
| Group 1 - 0 mg/kg | Mean | 5.713 | 0.650 | 4.853 | 0.130 | 0.050 | 0.013 |
| | SD | 2.827 | 0.202 | 2.542 | 0.046 | 0.036 | 0.006 |
| | N | 3 | 3 | 3 | 3 | 3 | 3 |
| Group 2 - 10 mg/kg | Mean | 5.910 | 0.910 | 4.733 | 0.180 | 0.043 | 0.010 |
| | SD | 0.262 | 0.295 | 0.123 | 0.030 | 0.015 | 0.000 |
| | N | 3 | 3 | 3 | 3 | 3 | 3 |
| Group 3 - 100 mg/kg | Mean | 7.010 | 1.037 | 5.633 | 0.213 | 0.063 | 0.017 |
| | SD | 0.972 | 0.159 | 0.886 | 0.122 | 0.012 | 0.012 |
| | N | 3 | 3 | 3 | 3 | 3 | 3 |
| Group 4 - 300 mg/kg | Mean | 6.247 | 0.963 | 5.073 | 0.130 | 0.047 | 0.010 |
| | SD | 1.619 | 0.172 | 1.492 | 0.040 | 0.015 | 0.000 |
| | N | 3 | 3 | 3 | 3 | 3 | 3 |

C. Mean Serum Chemistry Parameters

| Sex: Male | | NA (mmol/L) [g] | K (mmol/L) [g] | CL (mmol/L) [g] | CA (mg/dL) [g] | PHOS (mg/dL) [g] | BUN (mg/dL) [g] | CREA (mg/dL) [g] |
|---|---|---|---|---|---|---|---|---|
| Group 1 - 0 mg/kg | Mean | 141.7 | 5.47 | 106.3 | 10.07 | 9.10 | 11.3 | 0.47 |
| | SD | 1.5 | 0.35 | 1.5 | 0.06 | 0.20 | 1.5 | 0.15 |
| | N | 3 | 3 | 3 | 3 | 3 | 3 | 3 |
| Group 2 - 10 mg/kg | Mean | 141.3 | 5.00 | 106.7 | 10.00 | 8.77 | 10.7 | 0.47 |
| | SD | 0.6 | 0.20 | 1.5 | 0.26 | 0.50 | 1.2 | 0.06 |
| | N | 3 | 3 | 3 | 3 | 3 | 3 | 3 |
| Group 3 - 100 mg/kg | Mean | 140.3 | 5.27 | 106.3 | 10.13 | 9.20 | 11.0 | 0.53 |
| | SD | 0.6 | 0.15 | 0.6 | 0.57 | 0.35 | 2.0 | 0.23 |
| | N | 3 | 3 | 3 | 3 | 3 | 3 | 3 |
| Group 4 - 300 mg/kg | Mean | 141.0 | 5.03 | 106.7 | 10.23 | 8.97 | 10.3 | 0.53 |
| | SD | 1.0 | 0.21 | 2.1 | 0.12 | 0.29 | 3.1 | 0.06 |
| | N | 3 | 3 | 3 | 3 | 3 | 3 | 3 |
